# Spatial and temporal diversity of positive selection on shared haplotypes at the *PSCA* locus among worldwide human populations

**DOI:** 10.1101/2022.05.19.492596

**Authors:** Risa L. Iwasaki, Yoko Satta

## Abstract

Selection on standing genetic variation is important for rapid local genetic adaptation when the environment changes. We report that, for the prostate stem cell antigen (*PSCA*) gene, different populations have different target haplotypes, even though haplotypes are not unique to specific populations. The C-C-A haplotype, whereby the first C is located at rs2294008 of *PSCA* and is a low risk allele for gastric cancer, has become a target of positive selection mainly in Asian populations. Conversely, the C-A-G haplotype carrying the same C allele has become a selection target in African populations. However, Asian and African populations share both haplotypes, consistent with the haplotype divergence time (170 kya) prior to out-of-Africa. The frequency of C-C-A/C-A-G is 0.344/0.278 in Asia and 0.209/0.416 in Africa. 2D-SFS analysis revealed that the extent of intra-allelic variability of the target haplotype is extremely small in each local population, suggesting that C-C-A or C-A-G is under ongoing hard sweeps in local populations. From the TMRCA of selected haplotypes, the estimated onset times of positive selection were recent (3-55 kya), concurrently with population subdivision from a common ancestor. Additionally, estimated selection coefficients from ABC analysis were up to ∼3%, similar to those at other loci under recent positive selection on standing genetic variation. Phylogeny of local populations and TMRCA of selected haplotypes revealed that spatial and temporal switching of positive selection targets is a unique and novel feature of ongoing positive selection at *PSCA*. This switching may reflect potential of rapid adaptability to distinct environments.

## Introduction

Selective sweeps can be classified into two cases, whereby the target allele is either single (classic hard sweep) or multiple (soft sweep) in a tested population (Rees et al. 2020). The former describes that a *de novo* mutation emerges and the allele soon after becomes advantageous. Many neutrality tests are designed to detect signatures of this type of classic hard sweep. In contrast, the latter, soft sweep, includes more complicated processes of positive selection. One of such processes includes a typical case where a *de novo* mutation emerges within a particular haplotype but is maintained under neutrality. Through genetic drift, the haplotype number increases and is then maintained as standing genetic variation. Environmental change may result in the onset of selection on multiple haplotypes. In humans, both *MCPH1* and *ADAM17* are such examples in East Asian populations (Satta et al. 2020). *MCPH1* is associated with brain size development (Evans et al. 2005); haplogroup D at *MCPH1* is composed of multiple haplotypes that carry the derived C allele at G37995C and is a selection target. Similarly, there are multiple target haplotypes at *ADAM17* (Satta et al. 2020). *ADAM17* is involved in the *NRG-ERBB1* signaling pathway, and variants in genes of this pathway are associated with the risk of schizophrenia and psychiatric phenotypes (Pickrell et al. 2009). Other cases of soft sweeps include repeated advantageous alleles that emerged from different genetic backgrounds (recurrent mutations), as well as gene conversion of adaptive mutations into a new genetic background (Jones and Wakeley 2008; Rees et al. 2020).

As one of the major causes of soft sweeps, standing genetic variation in a mother population is reported to contribute to rapid adaptation of a daughter population to a novel niche. In sticklebacks, neutral polymorphism of genes in a mother population living in seawater became adaptive when a daughter population colonized from seawater to freshwater. For example, polymorphisms containing adaptive alleles at both *Eda* (Colosimo et al. 2005) and *KITLG* genes (Miller et al. 2007) associated with light skin pigmentation contributed to the adaptation to freshwater. Similarly, copy number variation in the *FADS2* gene (Ishikawa et al. 2019), which encodes an enzyme that synthesizes DHA, affects the survival rate in freshwater. Interestingly, orthologous *KITLG* in humans is also associated with light skin pigmentation. Standing genetic variation at this locus in Africa contributes to parallel adaptation for vitamin D synthesis due to the reduced UV light in both Asia and Europe (Miller et al. 2007). The common functional haplotype in both populations becomes a target of recent positive selection.

Schrider et al. (2017) scanned the human genome using a machine learning method S/HIC for detecting loci under completed positive selection; they revealed that most loci under selection are classified into soft sweeps (including standing genetic variation) rather than classic hard sweeps. Although Schrider et al. did not examine each case in detail, we expect many unexplored loci under completed/ongoing soft sweeps and that such loci may be associated with rapid adaptation to new environments.

Another approach for detecting signals of both soft and hard sweeps is the two-dimensional site frequency spectrum (2D-SFS) method. Summary statistics (Fc, Gc0 and Lc0) in 2D-SFS (Fujito et al. 2018; Satta et al. 2020) evaluate the decay of intra-allelic variability (IAV) of haplotypes carrying a target site (D-group) due to selective sweep compared with that of haplotypes under neutrality (A-group). The 2D-SFS method can identify a target site among different populations and distinguish signals of ongoing selection from completed selection based on three summary statistics. Combined with target haplotype TMRCA (tD), we can estimate the onset times of positive selection and reconstruct the history of positive selection in individual populations.

Humans migrated from Africa to most corners of the Earth during a short period of at most 70 thousand years (Soares et al. 2012; Haber et al. 2019) and subsequently have genetically adapted to various novel environments, e.g. reduced UV light, low temperature, high altitude, novel pathogens and dietary changes. Many studies have been conducted to investigate associations between signatures of positive selection and causal selective pressures (reviewed in Rees et al. 2020). At some of these signatures on target loci, onset times of positive selection are inferred to be relatively recent (within ∼50 thousand years). In some cases, the same allele parallelly became a selection target in genetically distinct populations (Xue et al. 2006; Yang et al. 2018). We previously detected three shared haplotypes of the prostate stem cell antigen (*PSCA*) gene in two East Asian populations, Japanese in Tokyo (JPT) and Han Chinese in Beijing (CHB) (Iwasaki et al. 2020). Two haplotypes carry a C allele at rs2294008 (T/C) in the first position (C-C-A/C-A-G haplotype) and are associated with low risk of gastric cancer (The Study Group of Millennium Genome Project for Cancer 2008; Tanikawa et al. 2012), and the other haplotype carrying a T allele (T-C-A) is associated with high risk of gastric cancer. Among the three, C-C-A/C-A-G haplotypes were found to be under ongoing positive selection. A signal of ongoing hard selective sweep was detected in the C-C-A haplotype alone in JPT. In contrast, such signals were detected in both C-C-A and C-A-G haplotypes in CHB, which is suggestive of an ongoing soft selective sweep in CHB. The three haplotypes are shared not only between JPT and CHB, but are also found worldwide including African populations. This supports that the two selected haplotypes, C-C-A and C-A-G, diverged before out-of-Africa. Indeed, the divergence time of the two haplotypes is estimated to be 170 kya (thousand years ago) (Albers and McVean 2020). We hypothesize that either or both of these haplotypes are under selection in different subpopulations in the 1000 genomes project (1KGP). To test this hypothesis, we apply the 2D-SFS method and examine the extent of IAV and tD (TMRCA). Furthermore, using an ABC framework, we estimate the onset times of selection as well as selection coefficients. Based on these results, we discuss unique features of this selection.

## Materials and Methods

### Test populations

We retrieved variant call format (VCF) data of 26 subpopulations in five superpopulations (East Asian (EAS), European (EUR), South Asian (SAS), American (AMR), and African (AFR)) from 1KGP Phase 3 (The 1000 Genomes Project Consortium 2015). We followed each population code in the definition of 1KGP and used the GRCh37 genomic positions for each SNP. The ancestral/derived alleles are defined in 1KGP. Neutrality test with Two-Dimensional Site Frequency Spectrum (2D-SFS) and TMRCA of target haplotype We defined 2D-SFS with φi,j, which is the sum of the number of SNPs whose configuration is *i* derived alleles in the D-group and *j* derived alleles in the A-group (Fujito et al. 2018; Satta et al. 2020). We evaluated IAV with a particular frequency of target allele by using φi,j. This method has the advantage of being able to detect incomplete and complete selective sweeps. We defined the core region for detection of selection by 2D-SFS in the following steps: we calculated *r*^2^ with rs2294008 per SNP and the average *r*^2^ in each 1000 bp non-overlapped sliding window, and then determined the boundaries of the core region based on SNPs with *r*^2^ > 0.75. For one of the African subpopulations, MSL, the size of the core region determined by the above method was too short (2000 bp) for analysis; thus, we simply defined the boundaries of the core region using SNPs with *r*^2^ > 0.75 without calculating average *r*^2^.

We calculated Q-values with Benjamini-Hochberg procedure (Benjamini and Hochberg 1995) to archive FDR controls for each of the summary statistics of 2D-SFS (Fc, Gc0 and Lc0) and evaluated their significances (Q-value < 0.05). If there were more than one significant summary statistic, we defined this as a signal of positive selection. We further classified this signal into completed or ongoing: significant values in Gc0 and Lc0 but not in Fc were interpreted as completed positive selection based on Satta et al. (2020), and otherwise classified as ongoing positive selection. Additionally, in the cases where only one summary statistic showed a significant value, we further calculated a combined P-value (Brown 1975) for the three summary statistics, and a highly significant combined P-value (P < 0.01) was also defined as positive selection.

Null distributions of the three summary statistics were generated using ms (Hudson 2002) under neutrality with demographic models based on Schaffner et al. (2005). From the simulated data, we chose at least 1000 replicates of simulated/pseudo target sites that had similar frequency to the true/observed frequency of the target site.

Additionally, we estimated TMRCA (tD) of the target haplotype (D-group). The value of *u*tD is the average number of substitutions per lineage in the D-group that is L_ζ_0/m, where 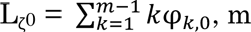 is the number of target alleles or haplotypes and *u* is the mutation rate per region per year (Satta et al. 2020). We applied 0.5 x 10^-9^/site/year for mutation rate (Scally and Durbin 2012) to calculate tD. For measuring tD in generations, the generation time is 29 years (Fenner 2005).

A framework of ABC for inferring onset of positive selection and selection coefficient (*s*)

We performed two simulation steps to obtain estimates of onset times of positive selection and selection coefficients (*s*) for three populations (JPT, YRI and CHB): (i) we generated samples using mssel implemented with positive selection (dominance coefficient (*h*) was set as 0.5) and accepted 10,000 replications with target site frequencies within ± 5% of observed target site frequency (Nakagome’s ABC procedure (Nakagome et al. 2019)); (ii) of these 10,000 replications, we accepted replications with summary statistics (F_c_, γ(10), γ*(10) and imax) that were within ± 1% of observed values in a test population. Finally, we estimated 95% CIs of *s* and onset times of positive selection from accepted replications.

C-C-A/C-A-G haplotypes are shared among all subpopulations studied (including African populations); this suggests that frequencies of these haplotypes should be higher than 1/2N, where N is effective population size of each local population. Therefore, we used a SSV (standing variant) model of mssel, but the uniform distribution (the prior distribution of allele frequency at selection onset (ft)) in the original model was modified with a simulated allele frequency distribution without recombination (see below).

We simulated the prior distribution of ft with the following procedure. We first defined the three haplotypes (C-C-A, C-A-G and T-C-A) based on three SNPs (rs2294008 (T/C), rs2976391 (C/A) and rs2978983 (A/G)). We then estimated the order of emergence of the three haplotypes based on derived/ancestral states and the divergence times of the three SNPs. The divergence times of the SNPs were estimated from the atlas of variant age (Albers and McVean 2020) (Fig. 1). C-C-A diverged from T-C-A 1,065,141 ya (36,729 gen. ago) as a *de novo* haplotype; then, C-A-G with two mutations (a derived A allele at rs2976391 (168,432 ya) and a derived G allele at rs2978983 (171,999 ya)) diverged from C-C-A 168,432 ya (5,808 gen. ago). Then, we simulated the frequency of a target haplotype for 10,000 replications, considering the above evolutionary process and demography of each test population based on Schaffner et al. (2005). The 10000 replications did not include cases where any haplotypes were fixed or lost before the onset of positive selection.

**Figure 1.**
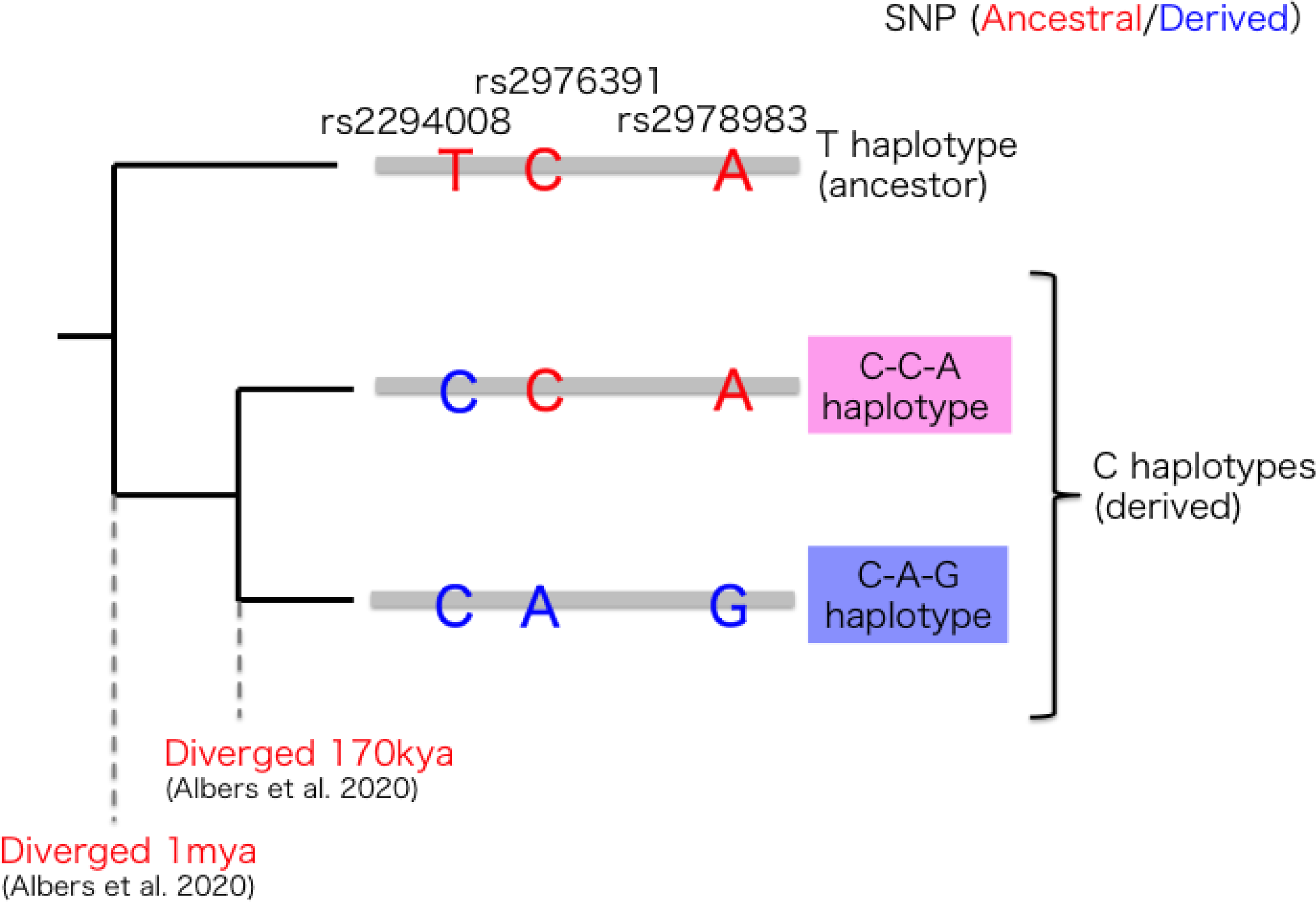
The relationships among haplotypes. The C haplotype, with a C allele at the first position of three SNPs (rs2294008, rs2976391 and rs2978983), diverged from the T haplotype around 1 mya (Albers and McVean 2020). Within the C haplotype, the C-A-G haplotype with two derived alleles at rs2976391 and rs2978983 diverged from the C-C-A haplotype around 170 kya (Albers and McVean 2020).

We generated a distribution of target allele frequency at the onset of positive selection respectively at ^tD^/_29_ generations ago. For JPT and YRI, we generated a distribution of each single target haplotype (C-C-A or C-A-G) frequency. In contrast, for CHB, we treated C-C-A and C-A-G as a single C haplotype. Then, the frequency of the C haplotype was set based on the summated frequencies of both haplotypes. We used tD of the C haplotype in CHB as 28,225 ya. We divided haplotype frequency distributions into bins from 0.0 to 1.0 with increments of 0.01 and drew a histogram as the ft prior distribution. Replications did not accept any case where ft exceeded observed target allele frequency in each test population (Nakagome et al. 2018).

## Results

### Different selection target among populations genetically closely related

Reduction in genetic diversity in the D-group (with core region strongly linked with a target allele) is due to selective sweep. This trait is exploited to detect signatures of positive selection on putative target alleles with the 2D-SFS method (Satta et al. 2020). Summary statistics of 2D-SFS are robust against violation of genetic diversity reduction caused by recombination (Iwasaki et al. 2020; Satta et al. 2020), and thus they can detect older signals of positive selection compared to haplotype-based tests (e.g. n*S*L) (Oleksyk et al. 2010; Ferrer-Admetlla et al. 2014). We applied the 2D-SFS method to 1KGP data to detect signals of positive selection on C-C-A and C-A-G haplotypes in different subpopulations.

No signals are detected on the two haplotypes in most of the AMR and EUR subpopulations (Table1). The frequency of the C-C-A haplotype is too low (<10%) in all subpopulations to detect signals. Especially, the C-C-A haplotype in both FIN and GBR show no private alleles in the D-group. Therefore, we conclude that all subpopulations in AMR and EUR have no signals of selection acting on the C-C-A haplotype, even though the combined P-value is marginally significant (e.g. GBR for P = 0.017). The C-A-G haplotype in PEL shows low values in Fc (Q = 0.002) and combined P (P = 0.016); PEL allele configuration shows two SNPs (rs75914363: φ87,0 = 1; rs2920288: φ92,0 = 1) that have similar number of derived alleles as the C-A-G haplotype (93 haplotypes). Consequently, this results in a low Fc value but high Gc0 and Lc0 values (Satta et al. 2020) and a significant combined P value.

Therefore, the presence of φ87,0 and φ92,0 is considered to be due to missampling of recombinants, φ87,j and φ92,j, where j is a positive integer. We conclude that the signal from PEL in C-A-G is false positive. The C-A-G haplotype in PUR and CEU show significant values in Lc0 alone, but the former shows a non-significant combined P value (P = 0.052); the latter shows a marginally significant combined P value (P = 0.046). Both signals in PUR and CEU can not be classified into any of the defined sweeps (Satta et al. 2020) (see Materials and Methods).

In non-AMR/EUR subpopulations, signals of selection are detected in either or both haplotypes. In all AFR subpopulations, ongoing positive selection on the C-A-G haplotype is detected.

Values of imax, γ(10) and γ*(10) are supportive of hard selective sweeps in AFR. Additionally, ESN and YRI show significant Lc0 values only in the C-C-A haplotype. ESN also shows a marginally significant combined P value (P = 0.040). For the same reason as in CEU, we do not regard this case as a positive selection signal. In contrast, YRI shows a non-significant combined P value (P = 0.062).

In SAS subpopulations, we observe three patterns of selection signals. First, a signal of ongoing positive selection on the C-C-A haplotype only in BEB is detected. Values of imax, γ(10) and γ*(10) in BEB are indicative/supportive of a hard selective sweep. Second, no signals for selection on any of the haplotypes in GIH and PJL are detected. Third, both STU and ITU show non-significant values of Fc but significant values of Gc0 and Lc0 in the C-C-A haplotype. This observation may imply that completed positive selection has acted on the C-C-A haplotype in both STU and ITU (see Materials and Methods).

In contrast, EAS subpopulations show two patterns of selection signals. We detect selection signals on C-C-A in JPT, CHS, KHV and CDX, and we find signals acting on both C-C-A and C-A-G haplotypes in CHB (Table 1). Based on the imax, γ(10) and γ*(10) values, these signals are all indicative of hard selective sweeps. For the C-A-G haplotype in CHS, there is only a significant Lc0 value (Q = 0.042) and a marginally significant combined P value (P = 0.032). For the same reasons as in CEU, we do not regard this as a positive selection signal. IAV of both haplotypes in CHB show signals of positive selection. Therefore, we conclude that the selective sweep acting on the two haplotypes in CHB appears to be soft. Variations in the pattern of selection signals are observed especially in Asian populations. This suggests that the target of selection can differ among subpopulations, even if they are genetically close and their habitat environments are seemingly similar.

**Table 1.**
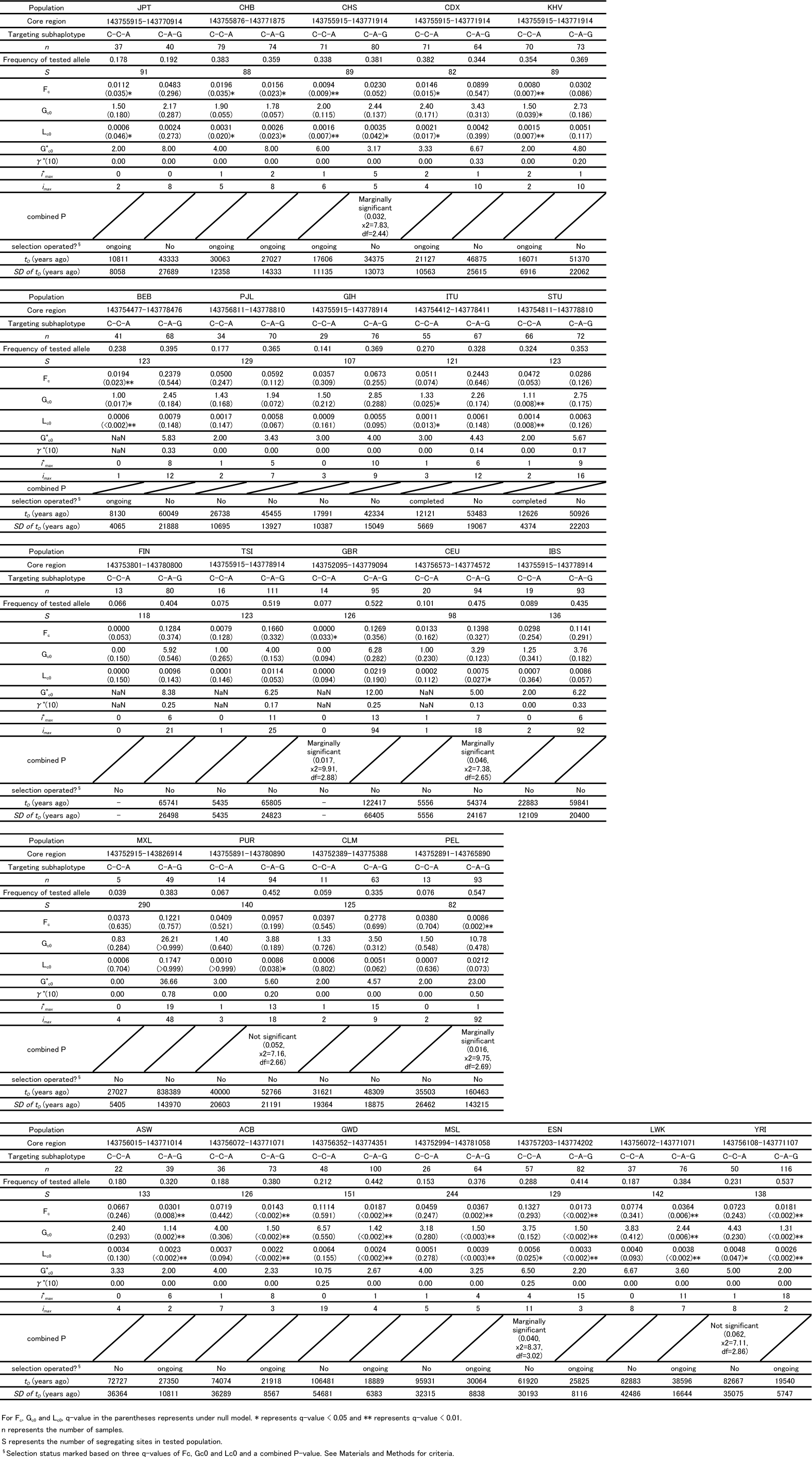
2D-SFS and tD values for 26 subpopulations.

### Inference of positive selection onset times

We estimate TMRCA of the D-group (tD ± SD) of the C-C-A or C-A-G haplotype (Fig. 2 and Table 1). TMRCA is the time for all D-group sequences to coalesce to a common ancestor (Satta et al. 2020). When the target haplotype is novel within a test population (hard selective sweep), the estimated tD is equivalent to the onset time of positive selection or earlier due to the waiting time of the *de novo* mutation (Satta et al. 2020). The values of tD of C-C-A haplotypes are limited within 8-30 kya (BEB: 8,130 ya for the minimum, CHB: 30,063 ya for the maximum) and those of C-A-G haplotypes are limited within 20-40 kya (YRI: 19,540 ya for the minimum, LWK: 38,596 ya for the maximum) in all subpopulations. The values of tD ± SD are limited within 3-42 kya for C-C-A and 13-55 kya for C-A-G. We also estimate the onset time for JPT/CHB/YRI using the ABC framework with samples generated with mssel (Nakagome et al. 2019) implemented with positive selection and compare them with the above tD values.

**Figure 2.**
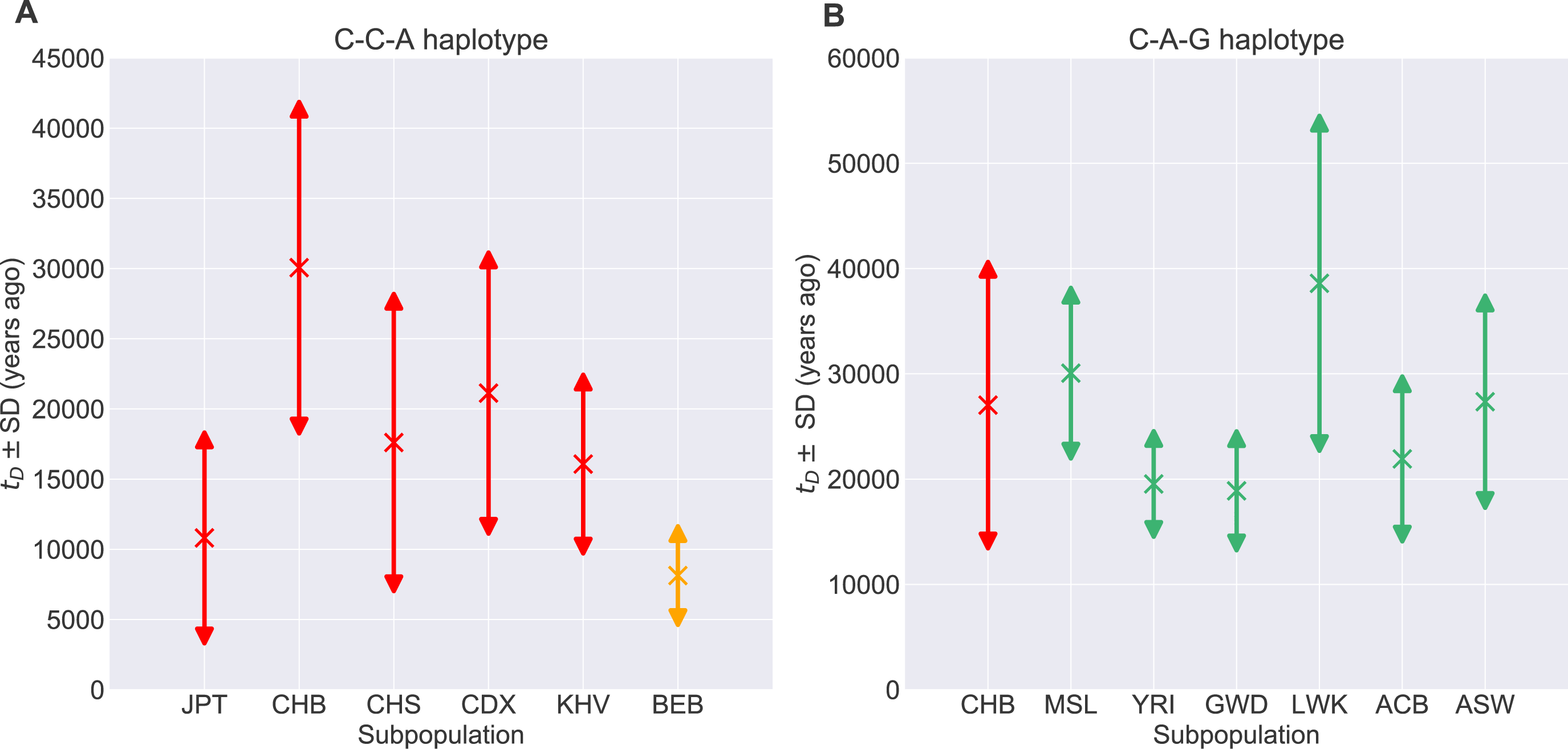
Values of tD ± SD in populations under positive selection are limited to within 3-55 kya. Note that the C-C-A haplotype in ESN and the C-A-G haplotype in CEU/PEL/CHS are excluded because they have marginal/unusual signals (see main text). (A) Values of tD ± SD for the C-C-A haplotype. (B) Values of tD ± SD for the C-A-G haplotype. The X markers represent the tD values and the double-sided arrows represent the ranges of tD ± SD. Red, orange and green represent East Asian, South Asian and African subpopulations, respectively

Mean onset times estimated by the ABC framework (ABC onsets) are 7,334 ya (C-C-A haplotype in JPT) and 13,845 ya (C-A-G haplotype in YRI). In CHB, two haplotypes have very similar tD values and they show significant low Fc and Lc0 values in 2D-SFS (Fig. 2 and Table 1); therefore, we regard both C-C-A and C-A-G as a single haplotype (C haplotype) carrying a derived (C) allele at rs2294008. The mean ABC onset in CHB is estimated to be 18,174 ya (Table 2 and Fig. 4A). All tD ± SD values are within the range of the 95% CI of ABC onsets.

**Table 2.** The onset times and selection coefficients inferred with ABC method.

| | Target haplotype | $s$ (mean) | $s$ (95% CI) | onset time<br>(mean (years)) | onset time<br>(95% CI (years)) |
| --- | --- | --- | --- | --- | --- |
| JPT | C-C-A | 0.0189 | 0.0011–0.0958 | 7334 | 1137–23270 |
| CHB | C haplotype | 0.0194 | 0.0018–0.0718 | 18174 | 3213–59276 |
| YRI | C-A-G | 0.0290 | 0.0029–0.1038 | 13845 | 2390–42027 |

Additionally, we infer selection coefficients using the ABC framework. Estimated means of *s* are 1.89% for C-C-A in JPT, 2.90% for C-A-G in YRI and 1.94% for the C haplotype in CHB; all *s* values are lower than 3% (Table 2 and Fig. 4B).

## Discussion

### Transition process of target haplotypes of positive selection

We estimate the number of transitions of target haplotypes along the human lineage in a parsimonious manner, considering tD values and current target haplotypes in subpopulations together with the phylogeny based on pairwise *F*ST values among 1KGP subpopulations (The 1000 Genomes Project Consortium 2015) (Fig. 3). We assume that positive selection did not act on any haplotypes in the human common ancestor (before out-of-Africa), as tD values of selected haplotypes are more recent than out-of-Africa. The estimation results suggest that positive selection on target haplotypes would cease/relax and operate several times within each subpopulation or lineage, leading to the extant subpopulations after population splits. For example, for EUR and AMR, although they possess the two haplotypes, no signatures of positive selection in *PSCA* are detected. In contrast, the C-A-G haplotype in African subpopulations has been selected independently four times (Fig. 3), and positive selection target switched between C-A-G and C-C-A haplotypes multiple times in Asian subpopulations (see below).

**Figure 3.**
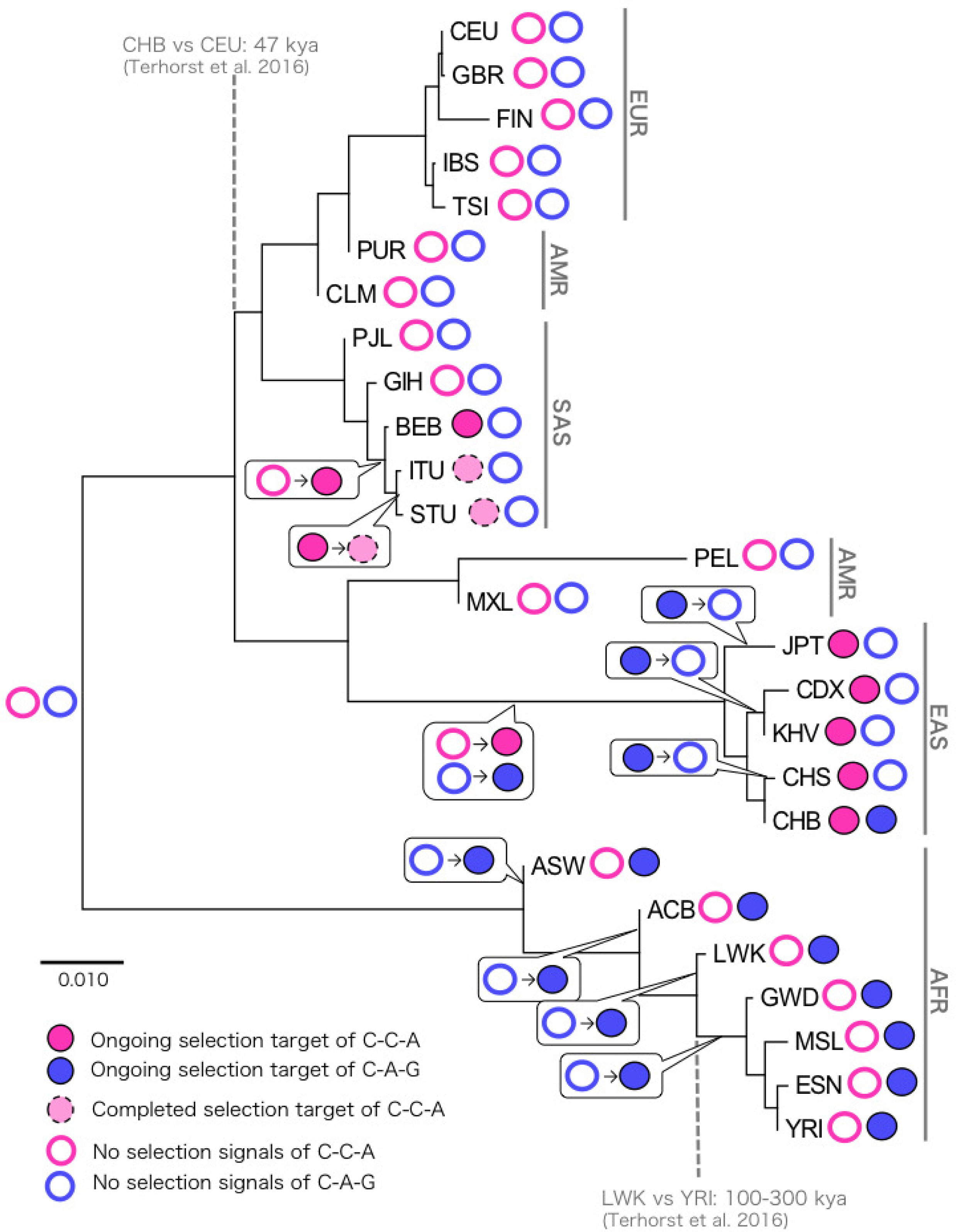
Transition of target alleles throughout human history. Summary of positive selection signals detected on C-A-G and C-C-A haplotypes in 1KGP 26 subpopulations. Filled pink/blue, dashed and open pink/blue circles indicate ongoing positive selection on C-C-A/C-A-G, completed positive selection on C-C-A and no significant signals of selection on C-C-A/C-A-G, respectively. The phylogeny of the 26 subpopulations was constructed using the neighbor-joining method of pairwise *F*ST values (The 1000 Genomes Project Consortium 2015). Transition process of selection target haplotype is inferred from the extant target haplotype, their tD values as well as subpopulation divergence times. In cases with no available divergence time data of subpopulations, we inferred timing of transition in a parsimonious manner.

In EAS subpopulations, target haplotypes are not the same between JPT and CHB, despite these populations being genetically close. The maximum tD of EAS is 30063 ya for the C-C-A haplotype and 27,027 ya for the C-A-G haplotype in CHB (Fig. 2). The Jomon population is one of the ancestral populations to the Japanese that split off from the stem lineage leading to the extant East Asian populations around 18-38 kya (Wang et al. 2018; Kanzawa-Kiriyama et al. 2019; Gakuhari et al. 2020). This period overlaps with the tD ± SD (13-42 kya) of the C-C-A and C-A-G haplotypes in CHB, suggesting that target haplotypes have been maintained from an East Asian common ancestor. In contrast, we do not detect any signals of positive selection on the C-A-G haplotype in JPT/CHS/CDX/KHV (Fig. 3 and Table1). This suggests that target haplotypes have transitioned from two haplotypes in a common ancestor (C-C-A and C-A-G) to a single haplotype (C-C-A) in all four populations. The phylogenetic relationship among CDX/KHV, CHS and JPT implies transitions are independent events in the three lineages.

Different from EAS subpopulations, only the C-C-A haplotype has been a target of selection in SAS. We detect a signal of ongoing positive selection in BEB and signals of completed positive selection in ITU and STU, whereas no selection signals are found in PJL and GIH. Parsimoniously considering these observations with the topology of the phylogeny (Fig. 3), selection on the C-C-A haplotype may have started to act in a common ancestor of BEB/ITU/STU. Then, positive selection was relaxed/ceased in a common ancestor of ITU and STU. In contrast, selection is still present in the extant BEB. The inferred divergence times of these SAS subpopulations have not been determined so far, thus making it difficult to discuss transition events of target haplotypes in detail.

In all AFR subpopulations, C-A-G is a common target haplotype. African demography is complex (Busby et al. 2016) and the examined African subpopulations share either or both western or/and eastern African ancestry with each other (Lachance et al. 2012; Excoffier et al. 2013; The 1000 Genomes Project Consortium 2015; Fan et al. 2019). One of the western African populations, YRI, and an admixed population of western African and eastern African, LWK (Henn et al. 2011; Excoffier et al. 2013; Gurdasani et al. 2015), diverged 100-300 kya (Terhorst et al. 2017). Although 1KGP did not cover subpopulations in eastern/southern Africa and there missing data of divergence times of some African subpopulations, the divergence time between LWK and YRI indicates that African subpopulations diverged earlier than the onset timings of positive selection (tD ± SD: 13-55 kya). The divergence time of the two subpopulations and the timing of positive selection parsimoniously suggest that positive selection on C-A-G in African subpopulations may have proceeded in parallel (Fig. 3).

Target haplotypes are different among subpopulations, revealing that adaptive alleles in each subpopulation are not always the same. Phylogenetic and tD analyses suggest that an adaptive allele in a subpopulation may change temporally and also differ spatially to other subpopulations. This change may follow some form of environmental transition. In both AFR subpopulations and CHB, the C-A-G haplotype is the target of selection and the values of tD are limited to more recent times, after out-of-Africa and subsequent population subdivision. This suggests that global environmental transitions may have caused a switch of selection operation among distinct subpopulations. In order to elucidate the global and local selective forces and to reconstruct the process of target haplotype transition, we need to survey selection signals in additional populations in the future. In particular, to determine the target haplotypes under selection in other populations and when target changed in other populations considering the EAS and Jomon split in the phylogeny, EAS subpopulations not included in 1KGP should be examined. Furthermore, we need to examine the association between global environmental change and the onset timing of positive selection.

### The relationships between tD and inferred onsets of positive selection

Using the approach by Satta et al. (2020), the timing of positive selection onset by calculating TMRCA of the D-group (tD ± SD) is estimated as 10,811 ± 8,058, 28,266 ± 9,300 and 19,540 ± 5,747 for C-C-A in JPT, the C haplotype in CHB and C-A-G in YRI, respectively (Fig. 2). We compare tD ± SD values with inferred onsets of positive selection from the ABC framework. Given the ft (allele frequency at selection onset) and current observed frequency, the ABC framework can estimate ABC onsets (i.e. the timing of positive selection operation) through simulations (Nakagome et al. 2019). As a result, for JPT, CHB and YRI, ranges of ABC onset 95% CI are wide enough to cover the respective tD ± SD values (Table 2 and Fig. 4A). We also compare tD ± SD with ABC onset 95% CI from Nakagome’s original ABC framework based on uniform distribution of ft (see Materials and Methods).

**Figure 4.**
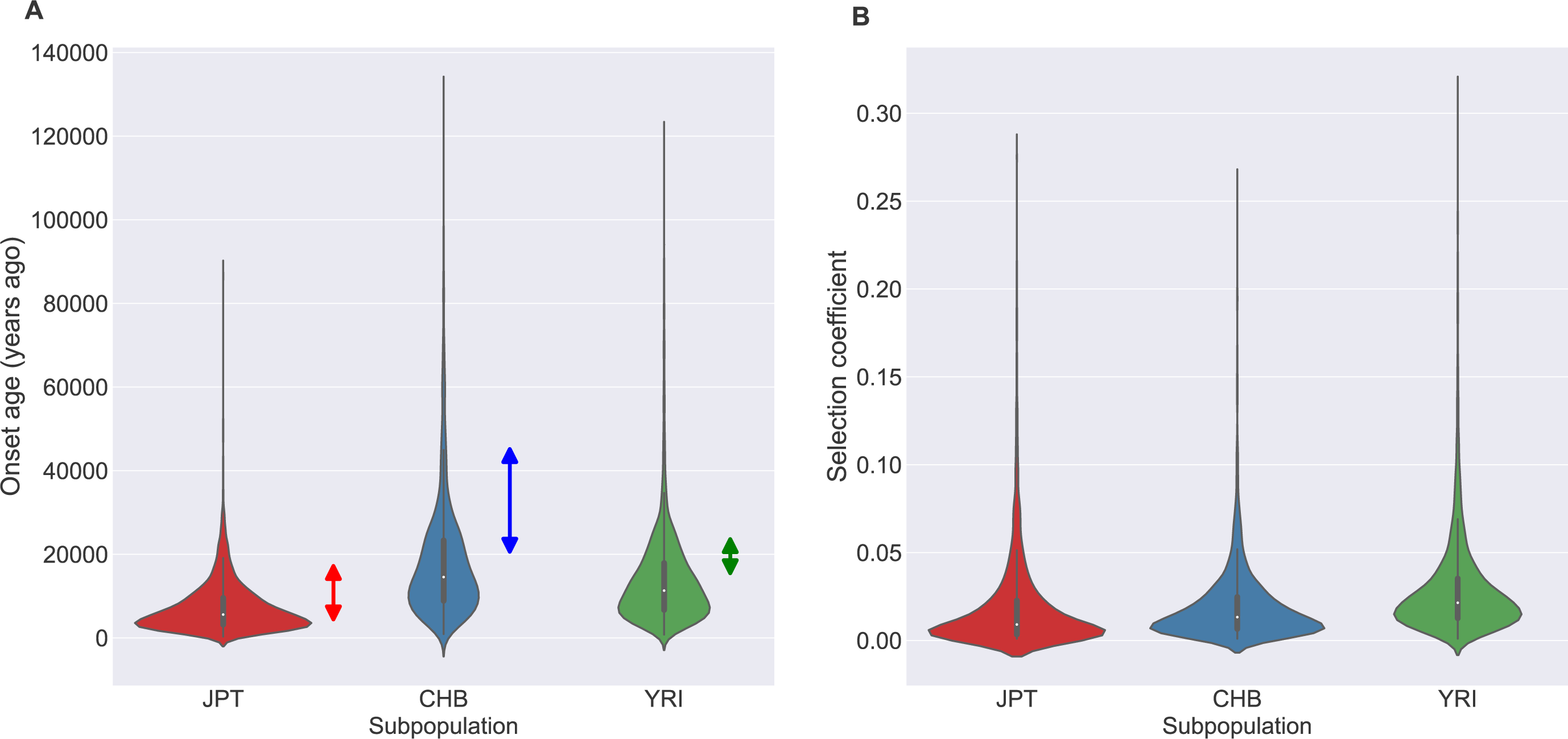
Inferred timing of onsets of positive selection and selection coefficients in JPT/CHB/YRI. (A) Violin plot of the timing of positive selection onsets estimated from the ABC framework. Double sided arrows indicate tD ± SD. See Table 2 for means (and 95% CI) of timing onsets of positive selection and Table 1 for ranges of tD ± SD. (B) Violin plot of selection coefficients. See Table 2 for means (and 95% CI) of selection coefficients.

Although ranges of the ABC onset 95% CI cover the respective tD ± SD values (data not shown), difference between the range of ABC onset 95% CI and tD ± SD is reduced if we use a prior ft distribution that is generated from simulation of ft rather than a uniform distribution. It is concluded that tD ± SD is supported by ABC onset 95% CI regardless of prior distributions of ft, suggesting that the ranges of tD ± SD are robust.

### Comparison of selection coefficients with those of other genes under selective sweeps

Selection coefficients (*s*) of the two haplotypes range from 1.9-2.9% in JPT/CHB/YRI (Table2). Since the tD of both haplotypes implies that positive selection likely acted recently, we first compare these *s* values to that of two representative genes under recent positive selection characterized to be involved in gene-culture coevolution: *LCT*, which is associated with lactase persistence among people of European ancestry (Gerbault et al. 2009; Itan et al. 2009; Peter et al. 2012), and *ADH1B*, which is associated with alcohol metabolism in CHB (Peter et al. 2012). The *s* of the C-C-A/C-A-G haplotypes are found to be equivalent to or smaller than *s* of *LCT* (*s* ≤ 9.6%) and *ADH1B* (*s* = 3.6%).

Next, we compare the *s* values with those of seven representative genes involved in responses to the outside environment and related to population differentiation. For *TRPM8*, a gene where selection acts in parallel among non-African populations (Key et al. 2018) and is associated with an endogenous response to cold temperature, *s* is estimated to be ∼1.5%; this is equivalent to the *s* of the two haplotypes. For the genes associated with skin pigmentation or morphology, most of the *s* values are equivalent to those of the two haplotypes: 1-2% for *KITLG* independently in East Asian and European populations (Beleza et al. 2013), 2-3% for *TYRP1* in European populations (Beleza et al. 2013), 4-5% for *SLC45A2* in European populations (Beleza et al. 2013) and 2.9% for *ASPM* in European populations (Peter et al. 2012). In contrast, *s* values at *EDAR* (*s* ≤ 14%; associated with hair and teeth morphology in East Asian populations (Kamberov et al. 2013; Peter et al. 2013)) and *SLC24A5* (*s* = 8-16%; associated with light skin pigmentation in European populations (Beleza et al. 2013)) show much larger values than those of the two haplotypes.

Finally, we assume that immune genes, which are associated with susceptibility to diseases, are under strong selective pressures, and thus, compare the *s* values of four representative immune genes with those of the two haplotypes. The *s* values of the two haplotypes are higher than most of those of *MHC* (*s* = 0.03-4%), which play a major role in immune responses (Yasukochi et al. 2013). A *CASP12* variant that is resistant to severe sepsis and under positive selection shows smaller values (*s* = 0.5-1% (Wang et al. 2006; Xue et al. 2006)) in human populations than that of the two haplotypes. In contrast, genes associated with resistance to malaria and under positive selection in African populations show larger *s* values (4.3% for *DARC* (McManus et al. 2017) and 20% > *s* > 10% (Saunders et al. 2005) or 4.4% (Tishkoff et al. 2001) for *G6PD*) than those of the two haplotypes. Therefore, the *s* values of the two haplotypes do not display conformation to any specific group of genes with particular function. Nevertheless, the *s* values of the two haplotypes (*s* = 1.9-2.9%), which emerged from standing genetic variation and under recent selection, are similar to *s* of another gene that also has haplotypes from standing genetic variation and under recent selection at the same time in distinct populations, *KITLG* (*s* = 1-2%) (Beleza et al. 2013).

### Comparison of onsets with those of other genes under selective sweep

Peter et al. (2012) assumed the C allele at rs2294008 as a single origin target allele and estimated onset age of selection and *s* in YRI. Their estimates for the onset age as 8000 ya (95%CI: 1000-54900) and *s* = 3.5% (95%CI: 0.4-15%) are in concordance with our estimations of the C-A-G haplotype in YRI. Smith et al. (2017) also estimated the onset age of selection of target alleles at five genes that are under recent ongoing positive selection but are not fixed in any populations of 1KGP. The study concluded that positive selection has acted in most human populations 50 kya or more recently. The values of tD ± SD (3-55 kya) for the selected C-C-A/C-A-G haplotypes overlap with the range of estimations by Smith et al. (2017). This suggests that the two haplotypes are also associated with recent differentiation among human populations.

### Temporal and spatial switching of target haplotype at *PSCA*

We consider that both C-C-A and C-A-G haplotypes each have a single origin and then spread by human expansion. The two haplotypes are locally under hard sweep in most subpopulations in EAS/SAS and AFR or soft sweep in CHB, although they are shared among all subpopulations tested.

Standing genetic variation contributes to rapid adaptation to environmental changes (Colosimo et al. 2005; Miller et al. 2007; Ishikawa et al. 2019). In *PSCA,* the two haplotypes are shared among all subpopulations tested. The observed major target allele in AFR is the C-A-G haplotype, whereas that in EAS/SAS is the C-C-A haplotype. Haplotypes derived from standing genetic variation respond differently and rapidly to the environments that each population encounters. This suggests that a particular local haplotype is not always adaptive among global human populations and such local adaptive haplotype(s) may not always be the same or only adaptive haplotype(s) in other subpopulations.

We also observe that target alleles can change within a lineage. For example, a target allele was both C-C-A and C-A-G in a common ancestor of EAS, but it changed to only C-C-A in JPT/CHS/CDX/KHV. This suggests that local ‘adaptive’ alleles can differ temporally and spatially.

In general, the same alleles emerged from standing genetic variation (e.g. *KITLG* in European/East Asian populations (Yang et al. 2018)) or independent *de novo* adaptive alleles in distinct populations (e.g. *LCT* in African/European populations (Tishkoff et al. 2007; Schlebusch et al. 2013)) are positively selected in many cases. In such cases, the selective force may be common among populations. We report that target haplotypes, C-C-A and C-A-G, switched temporally and spatially. Switching of target haplotypes may occur in future adaptations; therefore, these haplotypes may contribute as genetic sources of adaptation. Such target allele switching would provide a chance for individuals in a population to respond to a rapidly changing environment. We unveiled a unique and novel feature of positive selection in the *PSCA* locus and this study contributes to uncovering genetic diversity from the viewpoint of selection targets.

## Acknowledgements

The authors would like to thank Drs. Shigeki Nakagome, Naoki Osada, Hie Lim Kim, Tatsuya Ota and Jun Gojobori for valuable advice and discussion in this project, as well as colleagues in our University. Furthermore, we also thank Dr. Quintin Lau for English editing. This work has been supported in part by the Graduate University for Advanced Studies, SOKENDAI.

## Author contribution statement

RLI was responsible for reviewing the literature, conducting the research, extracting and analyzing data, interpreting results and writing the manuscript. YS was responsible for extracting and analyzing data, interpretation and discussion of results and amending the manuscript.

## Conflict of interest

The authors declare no conflict of interest.

## Notes

### Competing Interest Statement

The authors have declared no competing interest.

